# Transcriptional diversity in synaptic gene sets is sufficient to discriminate cortical neuronal identity

**DOI:** 10.1101/2022.11.25.517899

**Authors:** Amparo Roig Adam, Jose Martínez López, Sophie van der Spek, The SYNGO consortium, Patrick F Sullivan, August B. Smit, Matthijs Verhage, Jens Hjerling-Leffler

## Abstract

Synapse diversity has been described from different perspectives, ranging from the specific neurotransmitters released, to their diverse biophysical properties and proteome profiles. However, synapse diversity at the transcriptional level has not been systematically identified across all synapse populations in the brain. To quantify postsynaptic and identify specific synaptic features of neuronal cell types we combined the SynGO (Synaptic Gene Ontology) database with single-cell RNA sequencing data of the mouse neocortex. We show that cell types can be discriminated by synaptic genes alone with the same power as all genes. The cell type discriminatory power is not equally distributed across synaptic genes as we could identify functional categories and synaptic compartments with greater cell type specific expression. Synaptic genes, and specific SynGO categories, belonged to three different types of gene modules: gradient expression over all cell types, gradient expression in selected cell types and cell class- or type-specific profiles. This data provides a deeper understanding of synapse diversity in the neocortex and identifies potential markers to selectively identify synapses from specific neuronal populations.

## Introduction

Synapses are the information processing units of the brain and function in a use-dependent manner^1, 2^. Synapses are diverse with regards to subcellular targets and physiological properties^3^ and are central in information processing and storage theories^4^. Moreover, brain regions involved in higher cognitive functions, such as the hippocampus and neocortex, contain greater synapse diversity^5^. Synapse diversity also overlaps with the connectivity patterns (connectome) between brain areas associated to different functions^5^. Thus, understanding synapse diversity is crucial for gaining insights into the mechanisms for information processing in the brain.

Neurotransmitters and biophysical properties have traditionally been used for classification of synapses^2, 4^. New technologies, including diverse ‘*omics’*, are now uncovering additional layers of complexity and diversity on the molecular signatures of synapses^6^. Proteome differences between synapse types correlate with functional diversity (strength, kinetics, or synaptic plasticity)^4^. These different molecular profiles include, for example, scaffold proteins PSD95 and SAP102^5,7^, and AMPA-type glutamate receptors (AMPARs)^4^ for postsynaptic terminals of excitatory cells; and Gephyrin (GPHN) and Collybistin (ARHGEF9) scaffold proteins, and the GABAA receptors (GABAAR) for inhibitory postsynaptic sites^8^. On the presynaptic side, synaptotagmins 1 and 2, involved in calcium-dependent vesicle exocytosis, are differentially expressed between synapse types^9^. While most previous work is centered on inhibitory and excitatory synapses, recent studies have pointed out the expression of synaptic genes as correlated with neuronal diversity in subpopulations of transcriptomic cell types^10,11^.

Here, we aim to systematically identify how synaptic gene expression specifies the diversity of neuronal cell types in mouse neocortex using single cell transcriptomics data. Combining the expert-curated, evidence-based SynGO synaptic ontology^12^ and single cell expression data we could observe that expression of synaptic genes presents a striking diversity. Remarkably, specific biological function and synaptic component gene categories contained significantly high diversity and discrete modules of synaptic genes exhibited different modes of variability revealing that synapse diversity is organized at different levels.

## Methods

### Datasets

The single cell RNA-sequencing dataset published by Tasic et al. (2018)^13^ together with the information available in the SynGO^12^ database were retrieved for the study the expression of synaptic genes in the neocortex. The scRNA-seq dataset used was generated using the smart-seq RNA-sequencing technique. The tissue used was mouse primary visual cortex and anterior lateral motor cortex and up to 133 transcriptomic cell types and 16 cell classes were identified. The described cell classes include glutamatergic neurons, labeled according to their preferential layer of residence (for example Layer 4 neurons are labelled as L4) and their projection pattern (intratelencephalic, IT; pyramidal tract, PT; near-projecting, NP; and corticothalamic, CT) and GABAergic neurons labelled with their predominant expressed gene including: Sst, Pvalb, Vip, Lamp5, Sncg and Serpinf1^13^. The first release of the expert-curated synaptic gene ontology, SynGO1.0 was used^12^. SynGO1.0 contains 1112 unique human genes annotated to 2918 terms hierarchically organized and divided into ‘Cellular Components’ and ‘Biological Processes’ related to the synapse. These annotated synaptic genes encode evidenced proteins that localize to synaptic compartments and contribute to synaptic functions.

### Pre-processing and visualization

The scRNA-seq data were filtered to keep only the cells belonging to the classes ‘GABAergic’ and ‘Glutamatergic’ (n = 22439 cells) and expression data of the synaptic genes included in SynGO (1049 genes). Three subsets of the data were used for the downstream analysis separately by filtering the genes in the original dataset according to the following gene sets: all synaptic genes present in the SynGO database, SynGO presynaptic genes and SynGO postsynaptic genes.

Each of the filtered datasets was pre-processed with the standard protocol used by the Seurat^14^ R package as follows: log-normalization and scaling (scaling factor = 10000) of the raw count data, identification of highly variable genes, PCA dimensionality reduction, selection of significant principal components (PCs) by the Jackstraw procedure, and tSNE (t-distributed Stochastic Neighbor Embedding) dimensionality reduction/visualization (perplexity = 50). The significant PCs determined for each data subset were: 60 PCs for the full dataset, 42 PCs for the dataset with all synaptic genes and 21 PCs for both datasets with the pre- and postsynaptic genes. No quality filtering was performed on the cells since the dataset used was already passing the quality criteria in Tasic et al. (2018). The tSNE embedding of the data were color coded with the cluster identities determined by Tasic et al. (2018).

### Synaptic function and localisation (SynGO annotations) underlying diversity

MetaNeighbor^15^ was used to measure the power of each of the SynGO annotations to discriminate between different cell types. For each gene set (or SynGO annotated term), AUROC (Area Under the Receiver Operator Curve) scores were calculated for each of the 16 described cell types. To do so, random samples of the dataset were taken to train (2/3) the algorithm and test (1/3) the gene sets. Only those gene sets with at least 2 genes were used in this analysis. The result is an AUROC score for each gene set that can be interpreted as the performance of the gene set for the task of identifying each cell type, with 0.5 being equivalent to a random guess.

To calculate the statistical significance of the AUROC scores, the performance of random gene sets in MetaNeighbor was compared to that obtained with the SynGO annotations by generating random gene sets of the same size. For each gene set size in SynGO 10000 gene sets were generated by sampling from all the genes expressed in the original dataset, as well as all genes found in SynGO. For each of these randomly generated gene sets the ‘fast_version’ of MetaNeighbor was used. Firstly, the AUROC scores were used to compare the average performance of random gene sets and random synaptic gene sets of each size. Secondly, the random synaptic gene sets were used to calculate the statistical significance of the SynGO annotation scorings by calculating an empirical p-value. As indicated in Equation 1, this is done by calculating the fraction of scores from the randomized gene sets that are higher than the scores of the SynGO annotations. To calculate this as the overall score across all cell types, the score used for the p-value calculation was the sum of the AUROC scores in the 16 cell types. Likewise, we calculated the empirical p-value of SynGO annotations performing significantly worse than random gene sets (fraction of AUROC scores from randomized synaptic gene sets lower than the scores of SynGO annotations).

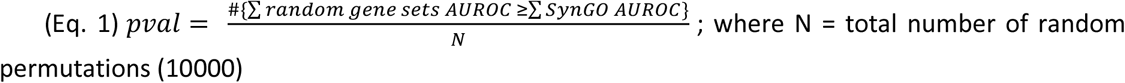

### Quantification of cell type diversity encoded by synaptic genes

To measure and compare the cell type diversity observed with different gene sets MetaNeighbor analysis was performed as described in the previous section. The quantified gene sets included the most high variable genes among: all genes in the dataset, non-synaptic genes (defined as all genes excluding the genes in SynGO), all synaptic genes, presynaptic genes, postsynaptic genes and mitochondrial genes (all genes included in the dataset and annotated in MitoCarta^18^). AUROC scores were calculated for each gene set and cell type, as well as for every cell class. Wilcoxon rank test (followed by false discovery rate [FDR] correction) was used to determine statistically different performance of each pair of gene sets.

### Synapse gene correlation network analysis

Weighted gene correlation network analysis (WCGNA) was used to investigate modules of synaptic genes in the transcriptional network of the dataset. In brief, the standard pipeline from the WGCNA^16^ R package was used to perform hierarchical clustering on the distance between every gene pair, calculated as 1-TOM (topological overlap matrix). To generate the TOM matrix, the co-expression similarity matrix is raised to a soft thresholding power (adjacency) that approximates a scale-free topology while keeping the mean connectivity of the network (ß = 4)^17^. Finally, the clustered genes are grouped into modules of highly interconnected genes using a dynamic branch-cutting algorithm. We used the dynamic tree cutting function (maximum height 0.9, 0.95, 0.98) and the modules were selected from a consensus of the result.

The gene modules were classified using K-means clustering on the eigenvector that explains the variance of gene expression in each cell type (80.3% variability explained). To do so, the average gene expression of each module in each cell was used to calculate the variance of expression for each gene module in each cluster. Next, the cell type identity information was removed, and the variance matrix ordered. The eigenvector explaining the maximum variability in the data (PC1) was used to cluster the modules in groups of similar variances of gene expression.

The individual gene modules were characterized using two approaches: mapping the average expression of the module to the synaptic types and annotating the function or cellular compartment they are related to by gene ontology enrichment. The former was done by mapping the average expression of all genes in each module, normalized to the average expression of random genes, to the transcriptomic cell types and visualizing it in the tSNE generated using only synaptic genes. To map the function and cellular component most related to each gene module, hypergeometric gene set enrichment was used. The background used for this analysis (universe) was comprised by all the genes annotated in SynGO that were present in the Tasic et al. (2018) dataset. The significance scores (p-value) from the hypergeometric tests were adjusted for multiple hypothesis testing using the Bonferroni correction method. Lastly, visualization of the test results for every gene module was produced with the sunburst custom color-coding tool of SynGO ontologies^12^.

### Code availability

https://github.com/Hjerling-Leffler-Lab/SynGO_scRNAseq

## Results

### Synapse genes contain cell identity information

To evaluate transcriptional diversity of synaptic genes in neuronal cell types, we filtered the expression data of cell types identified by Tasic et al. (2018) using the genes in SynGO. We analyzed four gene sets, including all genes in the original dataset (Fig 1A), all synaptic genes in SynGO (Fig 1B), presynaptic genes in SynGO (Fig 1C) and postsynaptic genes in SynGO (Fig 1D). We then compared the cell class and cell type diversity across the four different subsets. Here, we refer to the 133 transcriptomic neuronal types described in Tasic et al. (2018) as cell types (different colors in Fig 1) and 16 merged groups of these cell types as cell classes. Distinct classes and cell types could be discerned using SynGO genes only and all genes in the dataset to a similar extent (Fig 1A,B). Additionally, the observed transcriptomic diversity of presynaptic (Fig 1C) and postsynaptic (Fig 1D) genes showed similar levels of cell type specification. Quantification of the class and cell type discriminatory power of synaptic gene expression was calculated as the cell classification performance of each gene set using MetaNeighbor^15^. We included a similar sized gene-set from MitoCarta^18^ as comparison. Synaptic genes had a similar power in discerning classes and cell types in comparison to all genes or after removing SynGO genes (Fig 1E,F). We observed no difference in the discriminatory power between presynaptic and postsynaptic gene sets (Fig 1E, F). Similarly, no difference was observed when measuring pre- and post-synaptic genes in excitatory and inhibitory neurons independently (Suppl Fig 1A). However, there is a considerable overlap between the terms pre- and post-synaptic genes. Comparing only the genes specific for each category revealed a significantly higher score of postsynaptic genes for inhibitory neurons (Suppl Fig 1 B,C). This suggests that postsynaptic diversity is larger among GABAergic cells. These results indicate that the diversity of the synapse transcriptome across cell types is similar to that of the full transcriptome.

**Figure 1.**
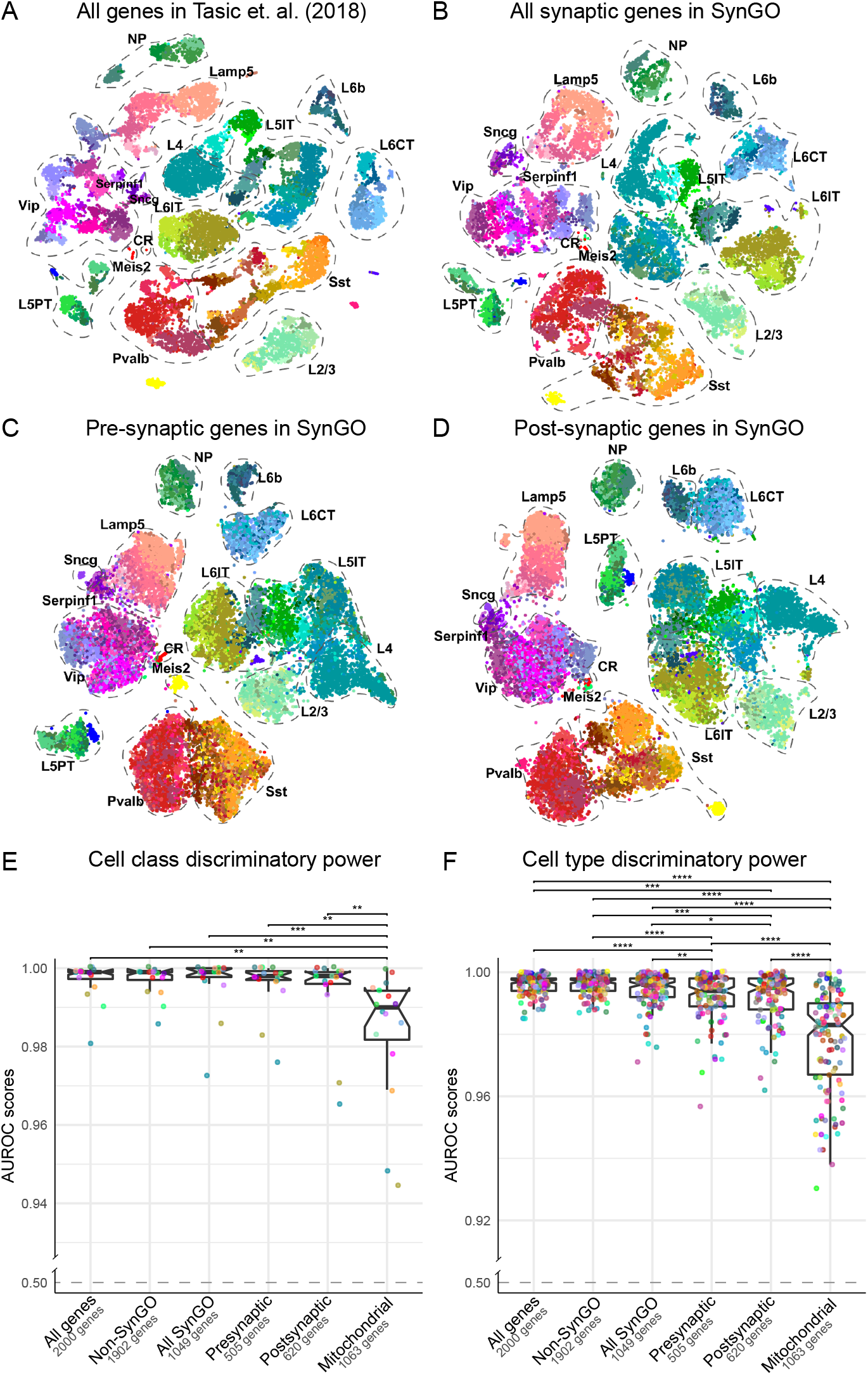
Neuronal diversity is recovered using only synaptic genes. t-SNE embedding visualization of the dataset from Tasic et.al. (2018) using the most variable genes among (A) all genes in the dataset, (B) only synaptic genes annotated in SynGO, (C) only presynaptic genes or (D) only postsynaptic genes, allow distinction of the annotated cell types to a similar extent. Dotted lines indicate cell classes and colors correspond to cell types described in Tasic et.al. (2018). Quantification was performed by calculating the cell class (E) and cell type (F) discriminatory power of each gene set in the MetaNeighbor pipeline. Wilcoxon rank test was used to determine statistically different performance of each pair of gene sets (*: p <= 0.05; **: p <= 0.01; ***: p <= 0.001; ****: p <= 0.0001).

### Specific SynGO annotations underlie synapse diversity

To identify whether genes contributing to synapse diversity belong to specific functional sets or are expressed in specific synaptic compartments, we analyzed the cell type discriminatory power of annotated SynGO terms. To test this, we used MetaNeighbor^15^ to score the performance of each SynGO term on the task of discriminating different cell types (AUROC scores) and compared it to random sets (of equivalent size) of genes drawn from SynGO and from all expressed genes in the dataset (Fig 2, Suppl Fig 2A). Several SynGO terms in both biological functions (BP, Fig 2B) and cellular components (CC, Fig 2C) discriminated cell types significantly better than random gene sets. Among the top biological process annotations are elements of the postsynaptic density organization, synaptic signaling, modulation of presynaptic chemical transmission and synaptic vesicle exocytosis. For cellular localisation, both presynaptic and postsynaptic membranes, as well as the presynaptic cytosol and active zone membrane were significant. A few categories conversely performed worse than random, including ribosomal genes and genes involved in metabolism (Suppl. Fig 2B,C). Analysis of average expression per category could not explain this result (Suppl Fig 2D,E). This analysis confirmed that synaptic genes perform better, on average, in cell type identification analysis than gene sets comprised of any gene expressed in the data also when normalising for number of genes. These results show that synapse diversity among different neuronal types accumulates in specific functions and cellular components.

**Figure 2.**
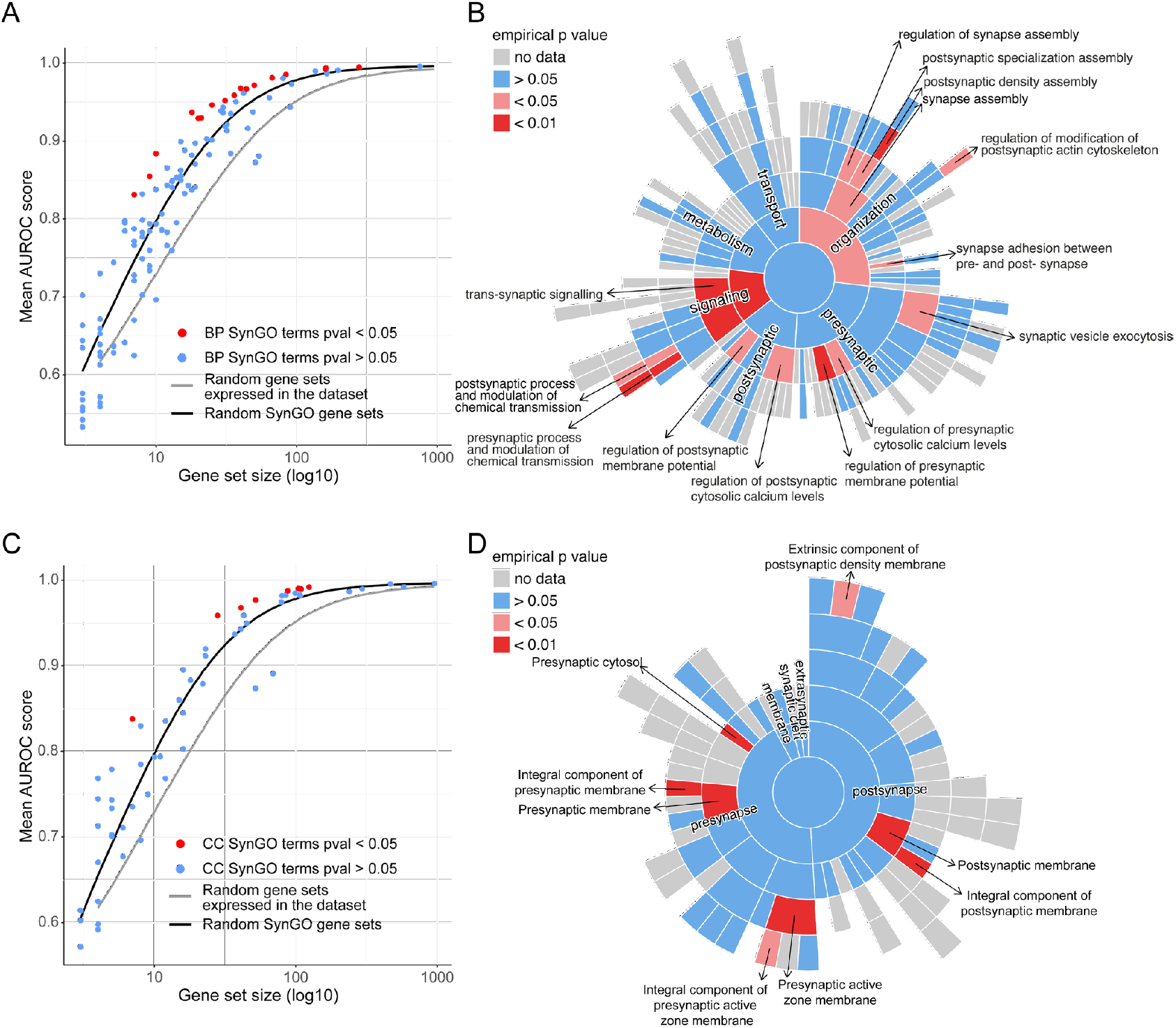
Synapse diversity resides in specific functions and cellular compartments. For all SynGO categories, the mean AUROC score across the 16 cell types is shown for biological functions (A) and cellular compartments (C) annotated in SynGO. Some SynGO terms (red) perform better than random synaptic gene sets of the same size (black line). Randomly generated synaptic gene sets (black line) discriminate cell types better than random gene sets (grey line) regardless the set size (Wilcoxon rank sum test; W = 3049, p = 0.001). The sunburst plots show the SynGO biological processes (B) and cellular compartments (D) where most variability lies across all neuronal subclasses. The color code (p-value) indicates SynGO terms that perform significantly better than random synaptic gene sets of the same size.

### Gene network analysis reveals different levels of synaptic organisation

WGCNA analysis and hierarchical clustering of the gene co-expression network revealed a high level of modularity of synaptic genes (Fig 3A). Classification of the gene modules according to the eigenvector calculated from the variance of gene expression across cell types (Fig. 3B), showed that synaptic gene modules can be clustered into three types: modules with specific expression in cell types or cell classes (discrete modules), modules showing a gradient of expression in a specific cell class (intermediate gradients) and modules with a similar gradient of expression in all cell types (pure gradients). Notably, these different classes of diversity were similarly found in pre- and postsynaptic gene modules. The results from this analysis were mapped to cell types using the average expression of the gene module in each cell (Fig 3E,G,I; Supp Fig 3). Interestingly, we observed modules with cell type specific expression in Vip-cells, sometimes shared with other cell types including Sncg-cells (pink; Fig 3I) and near projecting cells (dark green; Supp Fig 3). This suggest that some synaptic specializations can be re-used between GABAergic cell types and across GABAergic and excitatory cell types. In addition, gene set enrichment analysis of the obtained gene modules showed the biological processes and cell compartments (SynGO terms) to which each gene module is most related (Fig 3D,F,H). Interestingly, none of the enriched SynGO terms in the different groups of modules are overlapping between groups. Modules exhibiting gradients of expression included terms related to metabolism, post- and pre-synaptic ribosome, and protein translation (similar to those terms indicated in Suppl 2B). Interestingly we observed two gradient modules with opposing expression pattern (Fig 3C) suggesting that these are specific programs that are anti-regulated, perhaps in response to external signals or each other. This included the genes CTBP1 and ARL8 involved in “presynapse to nucleus signaling pathway” and “regulation of anterograde synaptic vesicle transport respectively” (yellow module), opposing the expression pattern of ribosomal and translational machinery genes (turquoise module). These results suggest that there are different types of synaptic organisation, ranging from cell type-specific to pan-neuronal programs, specified by distinct sets of genes at the transcriptome level, which also involve specific cellular functions.

**Figure 3.**
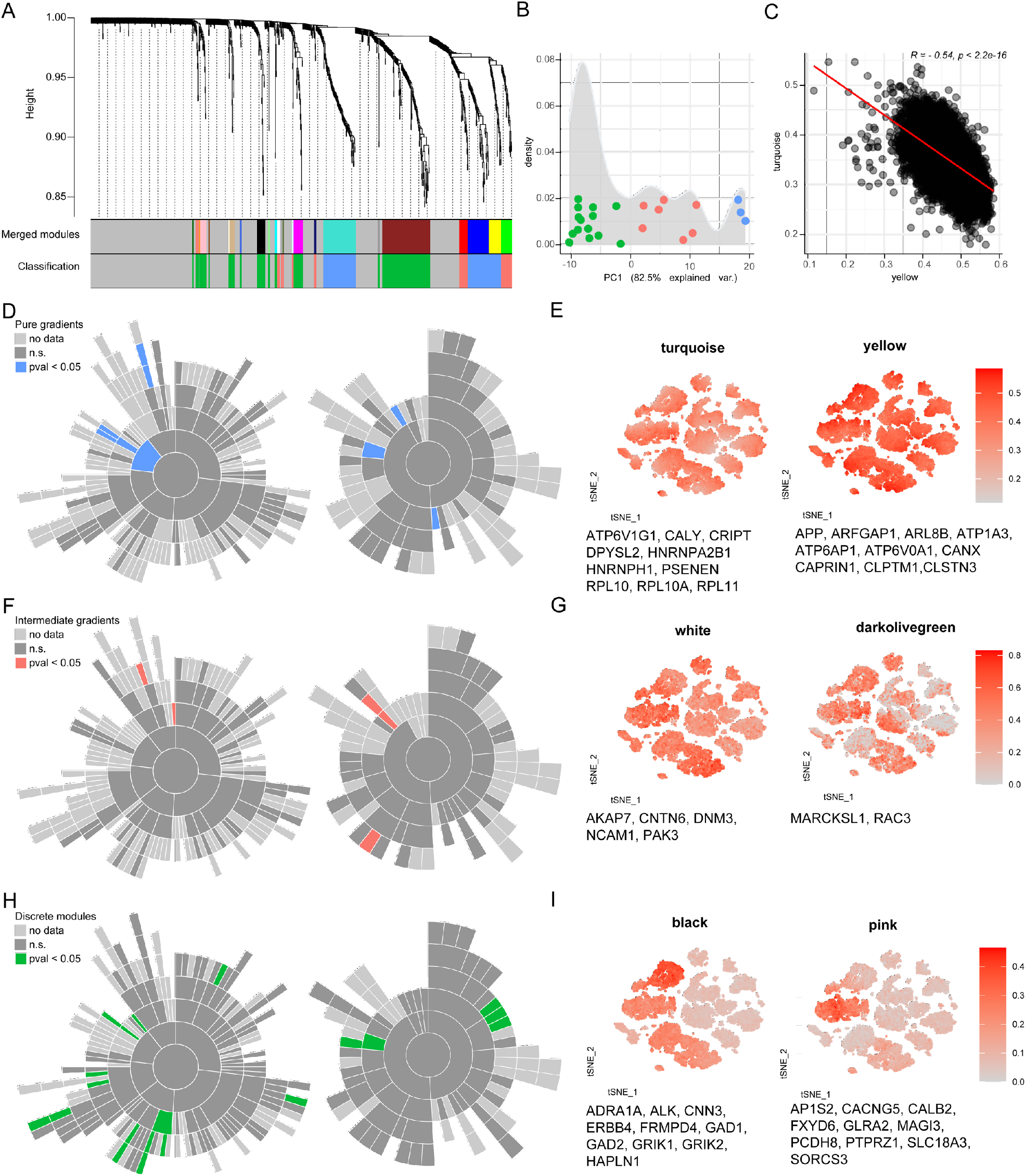
WGCNA reveals different levels of synaptic organisation: pure gradients within cell types, intermediate gradients and discrete expression in specific cell types and cell classes. (A) WGCNA dendrogram and gene modules selected. (B) Density distribution of the eigenvector (PC1) that explains the variance of gene expression across cell types in the gene modules (80.3% variance explained). Colour coding corresponds to the K-means gene module classification according to their greediness across cell types. (C) Anticorrelation of the average gene expression of the turquoise and yellow gradient modules. (D, F, H) Sunburst plots showing the biological functions (left) and cellular components (right) for which the gene modules of each type show enrichment. Dark grey indicates non-significant SynGO terms that contain genes in the modules, and coloured SynGO terms indicate enrichment in one of the gene modules. (E, G, I) tSNE plot of example modules within each group colour coded with the average gene expression of the genes in each module.

## Discussion

In this study, synapse diversity was mapped to previously defined transcriptomic neuronal cell types. We found that synaptic genes contain considerable cell identity information at the transcriptome level. Among synaptic genes, certain groups of genes associated to specific synaptic functions and localisation, annotated as SynGO terms, underlie the observed synapse diversity. Moreover, we identified additional candidate modules of co-expressed genes that contribute to synaptic functional diversity. These gene modules suggest different types of synapse organisation or different hierarchies of synaptic specification.

These results agree with the proposed vast synapse diversity arising from the combination of the different proteins that have been described as part of the synapse proteome^4,19^. Therefore, transcriptomic synapse diversity exists to a deeper extent of that depicted by the classical classifications of synapse types, possibly integrating the anatomical and physiological features classically described, as proposed for GABAergic interneurons in previous studies^11^.

We observed cell type-related diversity in both the pre- and postsynaptic genes. Our findings add additional gene-level resolution to the postsynaptic site diversity previously proposed in the brain based on protein expression of *Dlg4* (PSD95) and *Dlg3* (SAP102)^5^. Additionally, our data highlights the existence of such diversity also in the presynaptic site, showing a similar molecular diversity.

Our results show that synapse diversity, as well as similarity, between different cell types resides in specific synaptic functions and components. We identified cytoskeleton organisation, cell adhesion and synaptic signaling, as important for synapse diversity. As expected, we observed gene modules specific to excitatory/inhibitory synapse classification but also gene modules being specific to neuronal classes and neuronal types. An additional layer of diversity seems to be related to gradient-like expression of gene modules within each cell type, and surprisingly gene modules showing opposing expression which is likely an indication of dynamic synapse regulation as proposed by Zu F et al (2018)^5^.

Despite the single-neuron synapse diversity depicted here, recent studies have also described synapse diversity within a single neuron^6^. It is our hope that our results broaden the understanding of synapse diversity and generate hypothesis for future single synapse research. As an example, gene modules showing gradient expression profiles within cell types could reflect different cell states of the same cell types, in which single synapse variability could have a role. Our study provides the opportunity to expand the knowledge on the specific synaptic profile of distinct cell types. Further work in this direction could be used to selectively identify populations of synapses derived from specific populations of neuronal cell types, in intact tissue as well as in disease models.

## Acknowledgements

J.H.-L. was funded by the Swedish Research Council (Vetenskapsrådet, award 2018-00799), European Research Council (ERC) under the European Union’s Horizon 2020 research and innovation programme (Grant agreement No. 819540), and the Swedish Brain Foundation (Hjärnfonden). M.V., A.B.S. and A.R.A. were funded by the Broad Synapse 3 project (6910259-5500000759) and the Simons foundation SFARI director’s grant (882976).

## Notes

### Competing Interest Statement

The authors have declared no competing interest.

