## Supplementary Figures for "Transcriptional diversity in synaptic gene sets is sufficient to discriminate cortical neuronal identity"

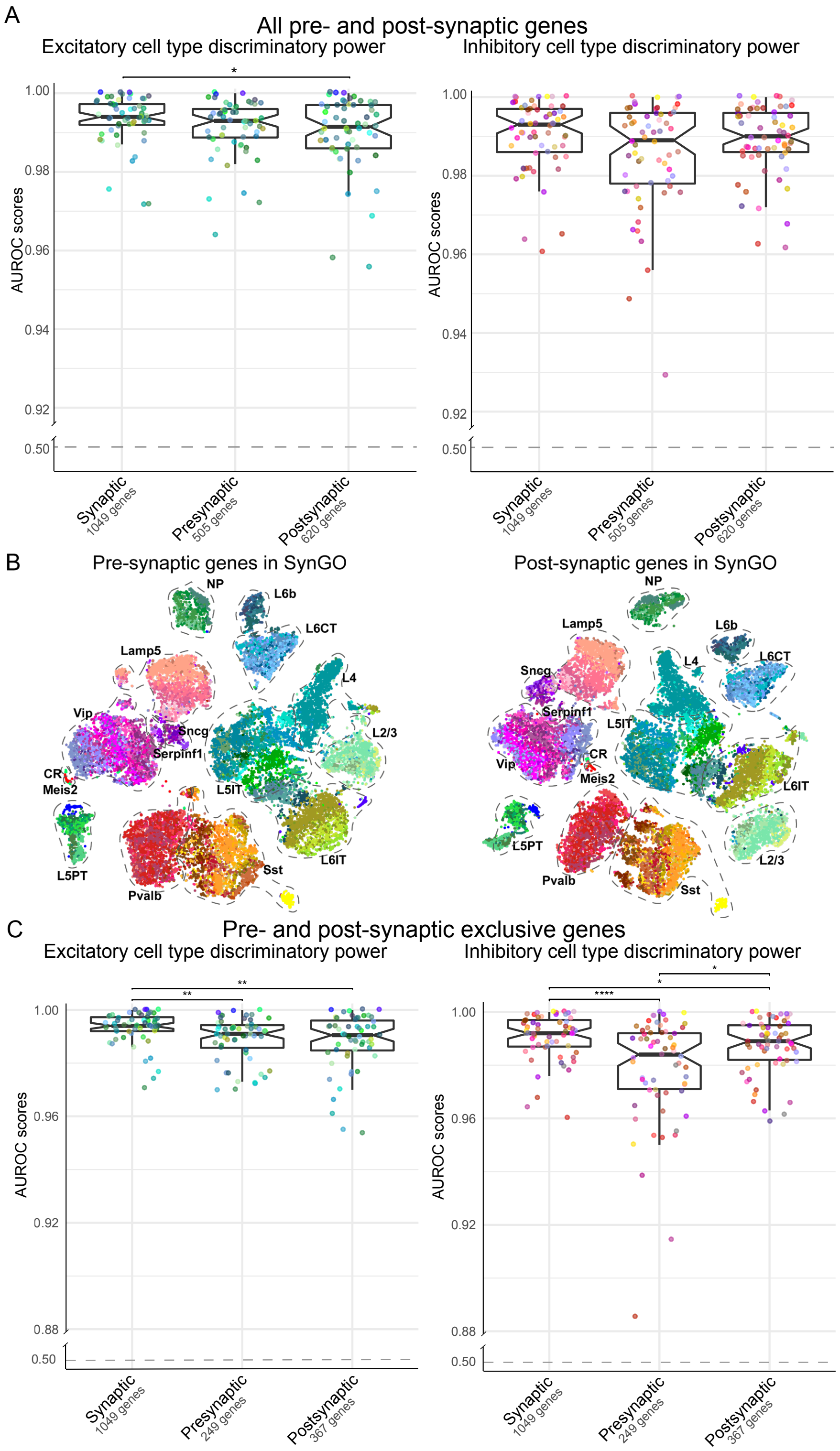


**Supplementary figure 1.** Comparison of cell type discriminatory power of pre and post synaptic genes within cell classes. Quantification cell type identity encoded in each all genes annotated in SynGO, all pre- and post-synaptic genes (A); and non-overlapping pre- and post-synaptic genes (C), as well as the tSNE embedding resulting from only exclusive pre- and post-synaptic genes are shown(C). Wilcoxon rank test was used to determine statistically different performance of each pair of gene sets (*: $p <= 0.05$; **: $p <= 0.01$; ***: $p <= 0.001$; ****: $p <= 0.0001$). Colour code of each cell type is the same used in Tasic et al. (2018) and Fig 1A.

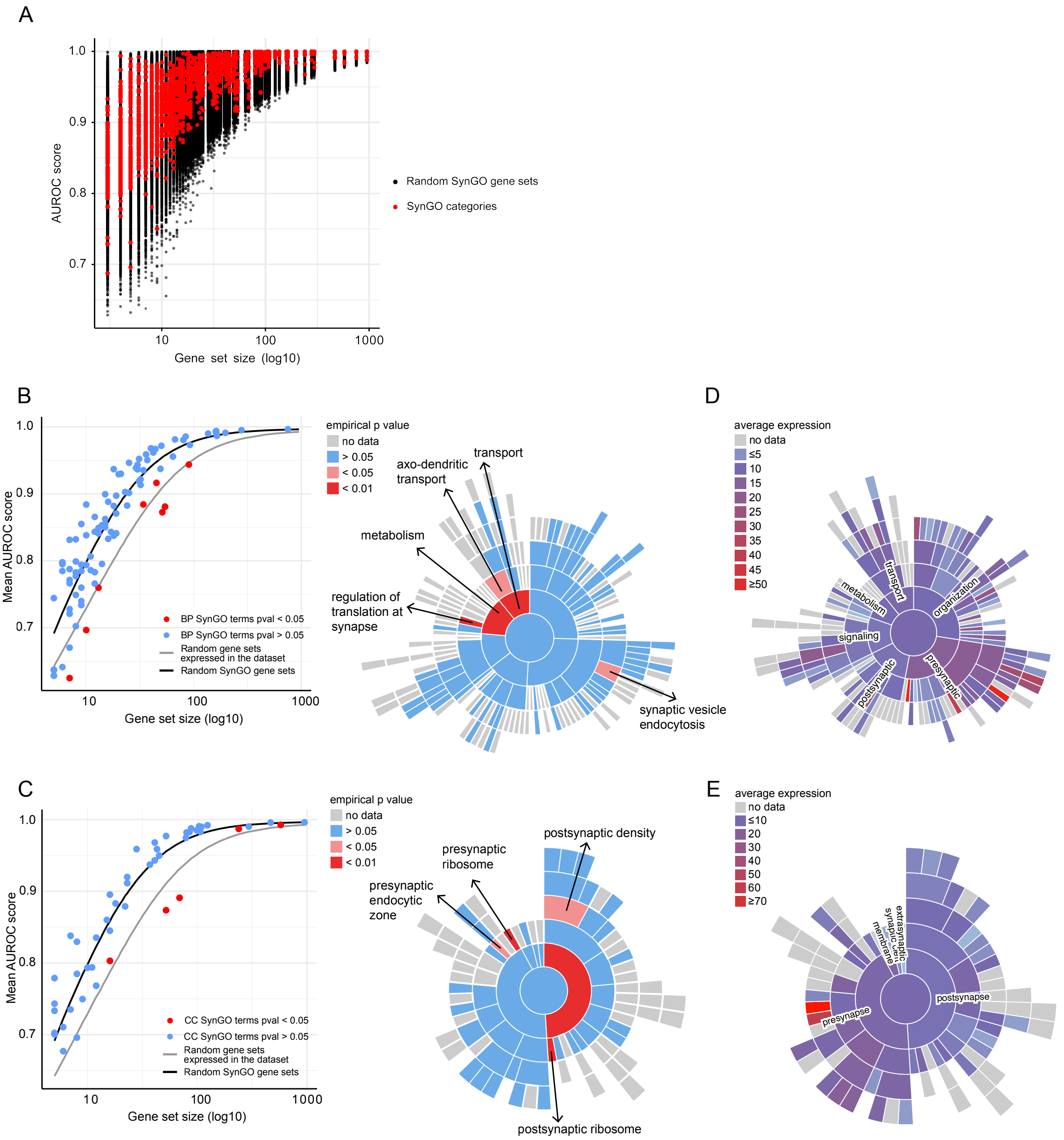


**Supplementary figure 2.** (A) All MetaNeighbor AUROC scores obtained in the random gene sets used for bootstrap analysis and annotated SynGO categories for each cell class in the dataset. (B and C) Mean AUROC score across the 16 cell types is shown for biological functions (B, left) and cellular compartments (C, left) annotated in SynGO. Some SynGO terms (red) score significantly worse than the random performance expected for their respective gene set size (black line). The sunburst plots show the SynGO biological processes (B, right) and cellular compartments (C, right) where less variability than expected lies across all neuronal subclasses. The colour code (p-value) indicates SynGO terms that perform significantly worse than random synaptic gene sets of the same size. (D and E) Average expression of all genes in each annotated SynGO term.

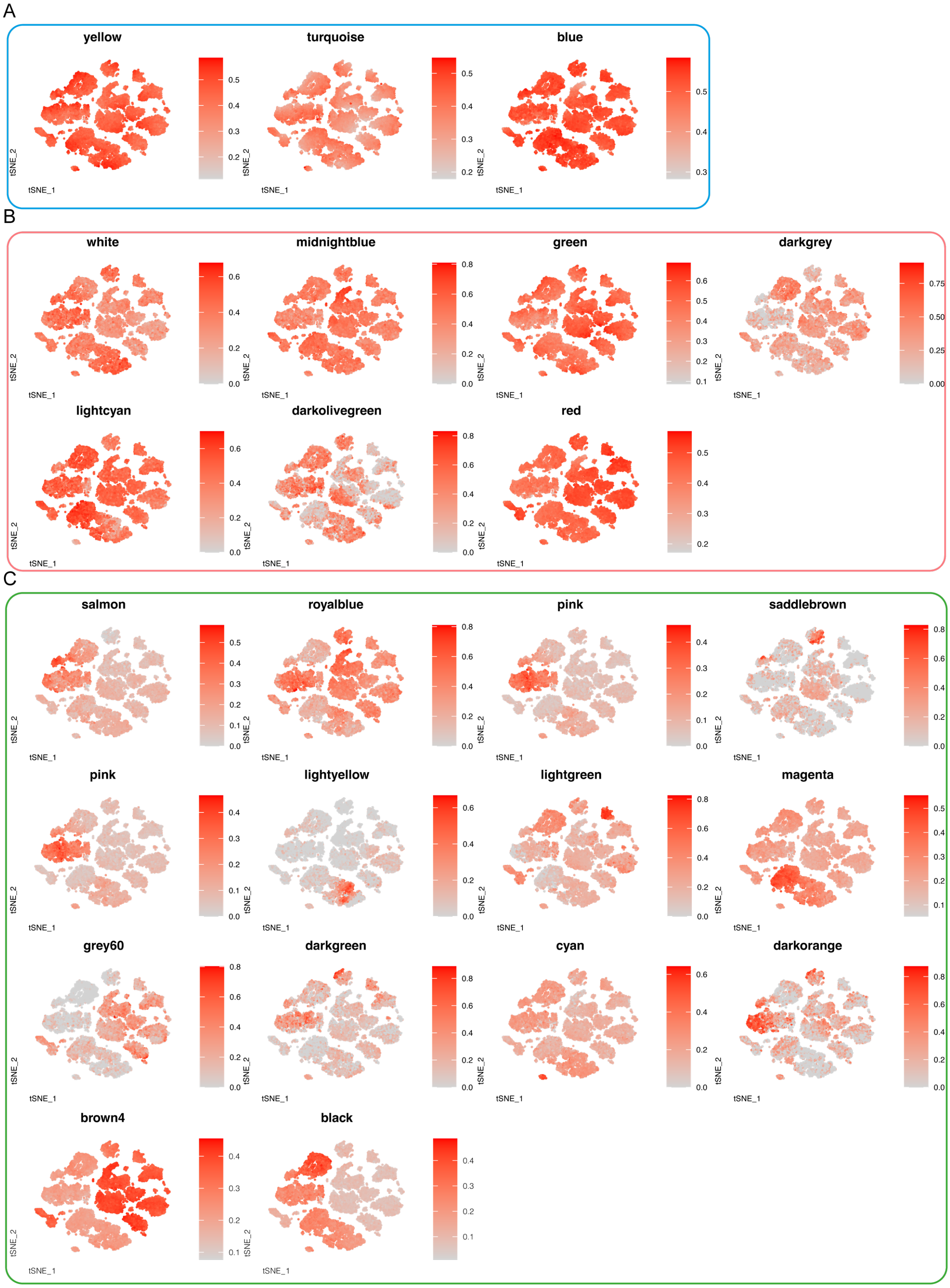


**Supplementary figure 3.** tSNE representation of the synaptic cell types (fig 1B) colour coded with the average expression of the genes in each module found with WGCNA.
